# Cell Fate Clusters in ICM Organoids Arise from Cell Fate Heredity & Division – a Modelling Approach

**DOI:** 10.1101/698928

**Authors:** Tim Liebisch, Armin Drusko, Biena Mathew, Ernst H. K. Stelzer, Sabine C. Fischer, Franziska Matthäus

## Abstract

During the mammalian preimplantation phase, cells undergo two subsequent cell fate decisions. During the first cell fate decision, cells become either part of an outer trophectoderm or part of the inner cell mass. Subsequently, the inner cell mass (ICM) segregates into an embryonic and an extraembryonic lineage, giving rise to the epiblast and the primitive endoderm, respectively. Inner cell mass organoids represent an experimental model system for preimplantation development, mimicking the second cell fate decision taking place in *in vivo* mouse embryos. In a previous study, the spatial pattern of the different cell lineage types was investigated. The study revealed that cells of the same fate tend to cluster stronger than expected for purely random cell fate decisions. Three major processes are hypothesised to contribute to the final cell fate arrangements at the mid and late blastocysts or 24 h old and 48 h old ICM organoids, respectively: 1) intra- and intercellular chemical signalling; 2) a cell sorting process; 3) cell proliferation. In order to quantify the influence of cell proliferation on the emergence of the observed cell lineage type clustering behaviour, we developed an agent-based model. Hereby, cells are mechanically interacting with direct neighbours, and exert adhesion and repulsion forces. The model was applied to compare several current assumptions of how inner cell mass neighbourhood structures are generated. We tested how different assumptions regarding cell fate switches affect the observed neighbourhood relationships. The model supports the hypothesis that initial cell fate acquisition is a stochastically driven process, taking place in the early development of inner cell mass organoids. The model further shows that the observed neighbourhood structures can emerge due to cell fate heredity during cell division and allows the inference of a time point for the cell fate decision.

**STATEMENT OF SIGNIFICANCE:** Cell fate decisions in early embryogenesis have been considered random events, causing a random cell fate distribution. Using an agent-based mathematical model, fitted to ICM organoid data, we show that the assumed random distribution of cell fates occurs only for a short time interval, as cell fate heredity and cell division quickly lead to spatial cell fate clustering. Our results show that neighbourhood clustering can emerge without specific neighbourhood interactions affecting the cell fate decision. The approach indicates four consecutive phases of early development: 1) co-expression of cell fate markers, 2) cell fate decision, 3) division and local cell fate clustering, and 4) phase separation, whereby only the phases 1-3 occur in ICM organoids during the first 24h of growth.

## INTRODUCTION

The first steps during mammalian embryo development are ovulation and fertilisation, followed by the preimplantation phase. At this point, the blastocyst is formed, which later implants into the uterus (1). Postimplantation development rapidly proceeds and involves multiple cell differentiation and morphological changes (1, 2). The first steps within the complex development processes in mammalian systems involve the cell fate decisions during the preimplantation phase. During development, the preimplantation phase is key to the success of pregnancy in mammals (3). Despite the importance of preimplantation, processes taking place during this phase are not fully understood.

The mouse is a common model organism to study the preimplantation phase. The 8 to 16 cells morula is formed until embryonic day 2.5 (E2.5) after fertilisation. At this stage, the first cell fate decision is taking place. Cells on the surface of the morula become trophectoderm (TE), while cells inside the morula are forming the inner cell mass (ICM) (4, 5). During E3.0 to E4.5, the second cell fate decision process takes place: ICM segregates into epiblast (Epi) and primitive endoderm (PrE) (6–8). NANOG and GATA6 are described as the first markers for Epi and PrE segregation, respectively. Expression levels of NANOG and GATA6 undergo progressive changes during the morula stage and the early blastocyst (9, 10). In early blastocysts (E3.0), all ICM cells seem to co-express NANOG and GATA6 (7, 11, 12). Subsequently, NANOG and GATA6 are gradually up- or down-regulated during the 32-cell stage. Thereby, both transcription factors repress each other locally (10, 13–17), leading to a mutually exclusive transcription factor expression in late blastocysts (64 cells) (18). Once a cell-fate is determined it is only possible to switch the fate by an external modulation of the included signalling pathways (16, 17, 19). While at early phases of preimplantation, the spatial distribution of NANOG and GATA6 positive cells is commonly described as a random salt-and-pepper pattern, GATA6 positive cells are sorted to the rim of the ICM in the late blastocyst and NANOG positive cells are forming an inner core (1, 7, 10, 11, 14).

Three major processes contribute to the final cell fate arrangements in mid and late blastocysts (reviewed in (20)). (1) Inter- and intracellular chemical signalling: the influence of NANOG, GATA6, and the FGF/ERK pathway onto the cell fate acquisition of single cells have been heavily studied over the past years (10, 13–15, 17, 19, 21–27). (2) Cell sorting processes: since cell lineage tracking revealed that ICM cells do not change their cell fate at E3.5, a cell sorting process is leading to the observed cell fate arrangements in late blastocyst (7, 10, 28–31). (3) Cell proliferation: from early blastocyst (E3.0) to late blastocyst (E4.5) it takes two to three rounds of cell divisions. Cell proliferation is hypothesised to contribute to the segregated state at E4.5 (32–34).

A recent study introduced a novel 3D stem cell system named *ICM organoids* (in the following the term *organoid* refers to the biological system, while the term *spheroid* is used in context of modelled data), which is based on inducible mouse embryonic stem cells (35). ICM organoids mimic the segregation into Epi and PrE without forming a TE and reproduce key events and timing of cell fate specification and cell-cycle progression in the ICM of mouse blastocysts. Thus, ICM organoids provide a powerful tool to develop and test biological preimplantation hypotheses *in vitro*. The cell fate of the cells within the ICM organoids are determined by their expression level of the transcription factors NANOG and GATA6. In total, four different cell types are described: mostly NANOG expressing cells which express a small amount of GATA6 (N_+_G_-_), mostly GATA6 expressing cells which express a small amount of NANOG (N_-_G_+_), cells that express NANOG and GATA6 on a high level (N_+_G_+_) and cells that express both transcription factors at a low level (N_-_G_-_). After 24 h of growth, most ICM organoid cells are either N_+_G_-_ or N_-_G_+_, meaning most ICM organoid cells are expressing one of both transcription factors at a high level and the other one at a low level, respectively. The spatial segregation into an inner core of N_+_G_-_ cells and an outer layer of N_-_G_+_ cells is visible after 48 h of growth. However, in contrast to mid mouse blastocysts, which consist of approximately 64 cells, ICM organoids comprise over 400 cells after 24 h of growth and more than 1000 cells after 48 h of growth. In order to quantify the patterns of neighbourhood distributions, a neighbourhood analysis of N_+_G_-_ cells, N_-_G_+_ cells, N_+_G_+_ cells and N_-_G_-_ cells was conducted in ICM organoids (35). Interestingly, the study revealed a local clustering of cells sharing the same expression type, inconsistent with the assumed salt-and-pepper pattern of cell fates.

The work of Mathew *et al.* (2019) (35) indicates that between the two stages of an initial salt-and-pepper distributed cell fate acquisition (E3.0-E3.5 early-blastocyst or 0 h old ICM organoids) and the final segregation of different cell fates (E4.5 late blastocyst or 48 h old ICM organoids), a local clustering of cell fates arises (E3.75 mid blastocyst or 24 h old ICM organoids). In order to test whether this pattern can be achieved through simple rules, static models were used. In these models, the cells are assigned to a cell fate but do not move. Different simple rules for these cell fate assignments were tested. However, none of the tested rules could reproduce the observed neighbourhood pattern of local clustering in 24 h old ICM organoids (35).

The purpose of this study is to investigate to which amount the observed neighbourhood structure in ICM organoids can be explained just considering cell divisions. To this end, a mathematical model is implemented. The implemented model is a 3D agent-based model. Agent-based models provide a technique to represent a wet-lab experiment under idealised conditions (36). Comparable agent-based models are commonly used to study cancer growth, cell proliferation or the contribution of single cells towards collective cell migrations (37–42). The model is given as a set of differential equations, describing mechanical cell-cell interactions, such as adhesion and repulsion forces, cell growth and cell division involving cell fate heredity. It is assumed that the initial cell fate acquisition results in a salt-and-pepper distribution at E3.5, which eventually leads to two segregated populations at E4.5 (10, 11, 29, 43). Hence, the initial cell fate decision is modelled as a stochastic process (omitting a detailed description of the signalling pathway dynamics) (44–46).

We use the model to investigate the hypothesis that the observed local cell fate clustering in 24 h old ICM organoids can arise from cell divisions alone, whereby cell fates are (partially) passed on to both daughter cells. Simulations were conducted under four hypotheses, each considering different cell fate switch rates during cell division. Our results indicate that the observed cell fate clustering can indeed be explained as a randomly distributed cell fate decision with subsequent divisions and cell fate heredity. Furthermore, based on the neighbourhood statistics, a time point for the cell fate decision (prior to the 24 h growth stage) can be inferred.

## MATERIALS AND METHODS

### Experimental data

Our study is based on experimental results from Mathew *et al.* (2019) (35). They used a system based on mouse embryonic stem cells which allows differentiation into primitive endoderm like cells (26). After induction of differentiation, 200 of these cells were placed in a nonadhesive well and centrifuged. Subsequently, the cells formed the 3D ICM organoids. This experimental approach implies that the cell fates in the initial organoid, composed of about 200 cells, are randomly distributed.

In 24 h and 48 h old ICM organoids, expression levels for NANOG and GATA6, as well as the nuclei positions for all cells were determined by fluorescence microscopy and subsequent image analysis. In total, four different expression types were established: NANOG and GATA6 expressing cells (double positive | N_+_G_+_), cells that express both transcription factors at a low level (double negative | N_-_G_-_), cells that express NANOG at a high cells and GATA6 at a low level (N_+_G_-_) and cells that express GATA6 at a high level and NANOG at a low level (N_-_G_+_).

In the following, the stage of 24 h old and 48 h old ICM organoids will be referred to as *t*_1_ and *t*_2_, respectively. During the simulations, cell fates and positions were recorded when the cell count in the simulation coincided with the average ICM organoid cell count in the experiment at *t*_1_ (441 cells at 24 h) and *t*_2_ (1041 cells at 48 h).

### Model implementation

An individual cell-based model that defines a small set of cellular features, implemented by Stichel *et al.* (2017) (42), is extended to explain the rise of local cell fate clustering in ICM organoids. The model describes the displacement of cells in response to external forces exerted by surrounding cells:

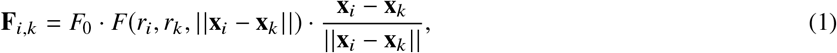

with *F*_0_ a positive constant, representing the strength of the mechanical interaction, *r*_*i*_ the radius of the *i*th cell, **x**_*i*_ = (*x*_*i*_, *y*_*i*_, *z*_*i*_) the position of the *i*th cell and

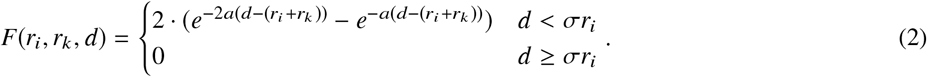

As given by the Morse potential (Eq. 2), the force between two cells is positive (repulsive), if the distance between the cell centres is below the sum of their radii, and negative (attractive) for (*r*_*i*_ + *r*_*k*_) < *d* < σ*r*_*i*_. Repulsion accounts for the limited compressibility of cells, while attraction accounts for cell-cell adhesion. The attractive part of the potential is cut off for cells at a distance above σ*r*_*i*_. If the distance of two cells equals the sum of their radii they are in perfect distance and thus neither exert attraction nor repulsion onto each other. The elasticity of the cells is given by the parameter *a*. The stiffness of cells increases with *a*. Displacement of cells in this model is only determined by forces exerted on them:

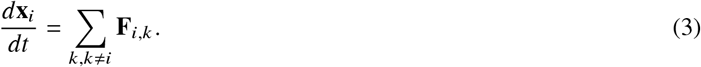

Each cell in this model is described by three features. A position **x**, a radius *r* and an expression type ∈. Other model parameters are assumed to be the same for all cells (e.g. elasticity, adhesion strength). The radius (size) of the cells is growing over time with

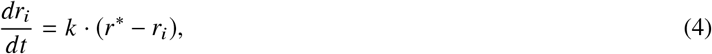

with *k* a (positive) growth constant and the maximum cell size *r*\*. Cell division is determined by a stochastic process which depends on the cell radius but not on the cell type. During cell division, the cell volume is preserved. The mother cell keeps its position (**x**_*m*_) and reduces its radius (*r*_*m*_) with:

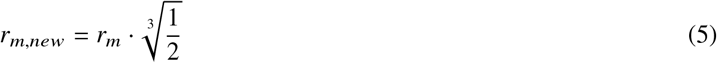

The daughter cell (**x**_*d*_) is generated close to **x**_*m*_, with **x**_*d*_ = **x**_*m*_ + *ξ* with *ξ* a random 3D vector (*δ*_*x*_, *δ*_*y*_, *δ*_*z*_) containing small values (*δ* ≪ *r*_*m,new*_). The daughter cell is assigned the same size as the mother cell (*r*_*d*_ = *r*_*m,new*_). The factor conserves the total cell volume during cell division. Directly after cell division, both cells are growing as given by Eq. 4 and change their positions as given by Eq. 3.

Since it was shown that the initial cell fate decision can be described as a stochastic process (44–46) and because the experimental approach leads to a random assembly of cell types, the initial expression type ∈ of the simulated cells is assigned randomly from the four expression types

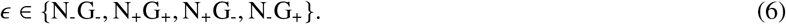

All parameters and the chosen values for the model are shown in Table A1.

### Cell fate heredity

The model can be used to test several assumptions addressing cell fate heredity. In total, four different hypotheses are tested. We consider the cell fate switches shown in Fig. A1 (i.e. N_-_G_-_ remain N_-_G_-_ or become N_+_G_+_; N_+_G_+_ remain N_+_G_+_ or become N_+_G_-_ or N_-_G_+_; N_+_G_-_ and N_-_G_+_ remain N_+_G_-_ and N_-_G_+_ or switch to the opposite cell fate).

Hypothesis 1 (H1) is considering no cell fate switches at all. The cell fate of a cell is strictly passed on to its daughter cells during cell division. Hypothesis 2 (H2) assumes that N_+_G_+_ produces N_+_G_+_ and N_-_G_+_ in equal parts during cell divisions, such that the amount of N_+_G_+_ remains constant over time, while the relative amount of N_-_G_+_ is increasing. We assume a third hypothesis (H3), with parameters such that the proportion transitions from *t*_1_ to *t*_2_ both are fitted. With a fourth hypothesis (H4) we consider a small cell fate transition rate from N_-_G_+_ to N_+_G_-_. The parameters and the chosen values for all hypotheses are shown in Tab. A2.

Please note that the fit involves only cell proportion data, and not the spatial positioning.

### Model framework

The implemented model is illustrated by a flowchart in Fig. A2. For all hypotheses, the initial cell fate proportion has been determined and assigned to occur at an earlier time point (*t*_0_). In particular, we couple *t*_0_ to the cell count and initiate the cell fate when the simulated ICM spheroid reached a cell count of 200, 300 or 400 cells. We determine the initial proportions of the cell fates by proportion data at *t*_1_ taken from the experimental data, omitting the spatial component. Using the system of linear differential equations (Eq. A1-A3), the theoretical proportions at *t*_0_ are determined. After cell fate acquisition, the simulation proceeds until *t*_1_ and *t*_2_ are reached. Both time points are coupled to cell counts as well (i.e. 441 cells at *t*_1_ and 1041 cells at *t*_2_). When *t*_1_ and *t*_2_ are reached, the model saves the spatially resolved cell positions and cell fates as output. Each simulation is repeated 100 times.

### Neighbourhood analysis and statistics

Cell neighbours were determined for both, simulation and experimental data, using the Delaunay triangulation. For the neighbourhood statistics, we derived the set of all neighbours of all cells of a given fate *j*, and computed the proportion of cell types based on this neighbourhood set. Neighbourhood proportions were collected for every executed simulation. For statistical comparison between experimental and simulated data according to single neighbourhood structures, we used the Wilcoxon-Mann-Whitney test with Bonferroni correction for multiple testing. In order to compare the overall fit of the simulated pattern to experimental data, we used the effect size as the relative deviation (ψ) as given in (35).

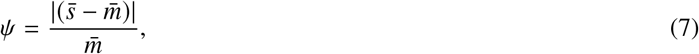

with 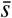 representing the mean of the simulated data and 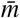 the mean of the experimental data.

### Spatial analysis of expression type distribution

During visual inspection of the biological data, we noticed that double negative cells often clustered at the rim of the ICM organoids. In order to visualise this effect, the following procedure was applied to experimental data for 24 h old and 48 h old ICM organoids. We assumed the spatial heterogeneity to be the same in small and in large ICM organoids, and therefore we normalised the size of all ICM organoids to equalise them in space. For normalisation, the median absolute divergence is used instead of the standard deviation because it is more robust with respect to outliers. The centre of mass for double negative cells was determined and the entire ICM organoid was rotated such that the centre of mass was located on the x-axis (i.e. *x* > 0, *y* = 0, *z* = 0). The rotated cell positions were combined in one data set. Each double negative cell was assigned the value 1 and the value 0 was assigned to all other cells. Interpolation on the combined data generated a continuous clustering pattern from experimental data. In the interpolated dataset, the label 1 indicated the presence of a particular expression type, while 0 indicated its absence. The procedure was repeated for double positive, NANOG positive and GATA6 positive cells to show that the visualised heterogeneity is not an artefact resulting of this procedure.

In order to analyse the cell fate proportions in dependence on their relative distance to the ICM organoid centre, we measured the distance of the cells of each ICM organoid to their respective centre of mass. Subsequently, we divided these distances into ten intervals. Eventually the mean proportions and standard deviations for the ten intervals were determined for all 24 h old ICM organoids and 48 h old ICM organoids. Unless otherwise stated, the model and data analysis methods were implemented in MATLAB R2019b.

## RESULTS

The purpose of the simulations was to study, if the complex 3D neighbourhood pattern in the ICM organoids can be explained by cell divisions alone. To this end, we combined several simple mechanisms, i.e. cell mechanics, a stochastic expression type assignment and cell fate heredity during cell division, but omitted cell type segregation processes, intra- and extracellular signalling.

We show that the simulations can be used to predict the ICM organoid cell count when the cell fate decision occurs. For this reason, we used a mechanistic mathematical model for 3D spheroid growth in order to investigate the distribution pattern of cell fates in 24 h old ICM organoids (see Methods). During the simulation, the cells were assigned to the different expression types (i.e. N_-_G_-_, N_+_G_+_, N_+_G_-_ and N_-_G_+_) randomly at *t*_0_, whereby the cell type proportions were set to fit the experimental data for 24 h old ICM organoids. The time point for the initial expression type assignment (*t*_0_) was coupled to cell count of the spheroid (200, 300 or 400 cells). One 24 h old ICM organoid, one 48 h old ICM organoid, as well as simulated ICM spheroids for *t*_1_ and *t*_2_ are shown in Fig. 1 A.

**Figure 1:**
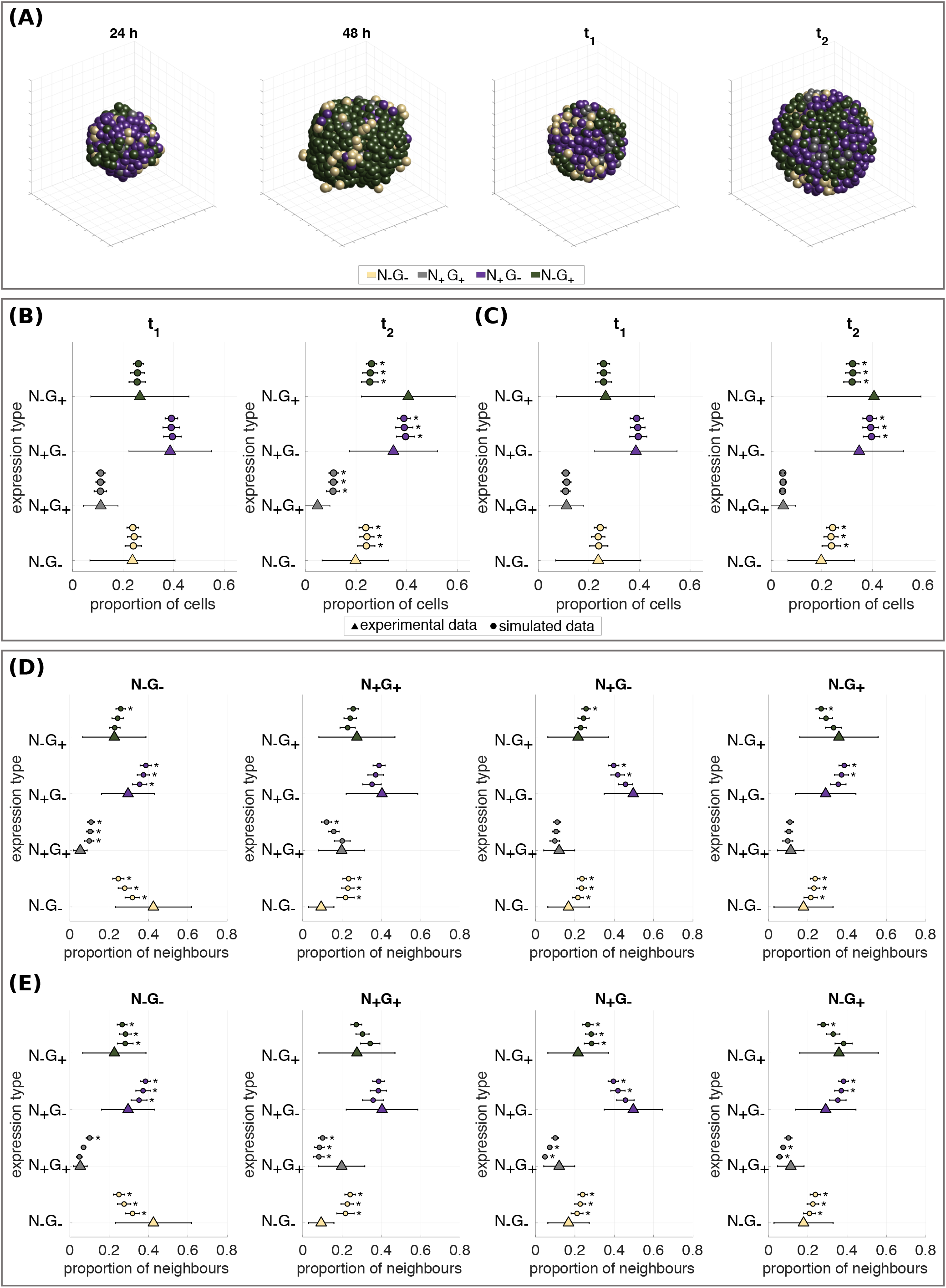
Expression type composition of ICM spheroids for H1 and H2. (A) ICM organoids (experimental data) for 24 h and 48 h and simulated ICM spheroids for *t*_1_ and *t*_2_. (B, C) Expression type composition of ICM organoids and ICM spheroids shown as percentage of the total number of cells within ICM organoids at *t*_1_ and *t*_2_. Statistically significant differences between the cell fate proportion of ICM organoids and ICM spheroids are indicated by stars (*p* < 0.05; using a Wilcoxon-Mann-Whitney test with Bonferroni correction). Simulations were performed under the assumption H1 (B) and the assumption H2 (C). (D, E) Expression type composition of neighbouring cells shown as percentage of the total of neighbouring cells at *t*_1_. Simulations were performed under the assumption H1 (D) and the assumption H2 (E). Experimental data from Mathew *et al.* (2019) are indicated by triangles. Simulation results for different *t*_0_ are indicated by circles. The error bars indicate the standard deviation. *t*_0_ from lowest line to top: 200, 300 and 400 cells. Statistically significant differences between the neighbourhood structure of 24 h old ICM organoids and ICM spheroid patterns are indicated by stars (*p* < 0.05; using a Wilcoxon-Mann-Whitney test with Bonferroni correction).

We tested four hypotheses addressing different cell fate heredity strategies (see Methods). In assumption H1, each cell passes on its cell fate to both daughter cells. For assumption H2, N_+_G_+_ are allowed to give rise to N_+_G_+_ or N_-_G_+_ cells. The production of N_-_G_+_ cells from N_+_G_+_ cells is sensible, because a) a switch from N_+_G_-_ to N_-_G_+_ or vice versa is biologically not feasible (16, 17), b) the amount of N_+_G_+_ cells remains constant between the time points *t*_1_ and *t*_2_, and c) the proportion of N_-_G_+_ cells increases strongly from *t*_1_ to *t*_2_. Although cell fate switches from N_+_G_-_ to N_-_G_+_ or vice versa are biologically not feasible we allow them in H3 and H4 in order to fit the proportion data measured at *t*_2_. H3 allows cell fate switches from N_+_G_-_ to N_-_G_+_ and H4 considers a small flux between N_+_G_-_ and N_-_G_+_ cells (see Fig. A1 and Tab. A2).

To determine the hypotheses with the closest fit to the experimental data, the effect size as the relative deviation of the simulated data is calculated (see Fig. A3). H1 and H2 show the closest fit to the experimental data (see Fig. 1). The results for H3 and H4 are shown in the supplement (see. Fig. A4).

Starting from the initial cell fate assignment at *t*_0_, cell growth, displacement according to adhesion and repulsion forces, and cell divisions were simulated according to Eqs. 1-6. Cell fates and positions were recorded at *t*_1_ and *t*_2_.

The expression type compositions of simulated ICM spheroids were determined and statistically compared with experimental data using the Wilcoxon-Mann-Whitney test with Bonferroni correction. For H1 (Fig. 1 B) and also for H2 (Fig. 1 C), the simulated proportions agreed very well with the experimental data at *t*_1_. At *t*_2_, the simulated proportion data showed for the assumption H1 significant differences to experimental data for all expression types at all cell fate initialisation time points *t*_0_ (*p* < 0.05). For the assumption H2 (Fig. 1 C), significant differences between predicted and experimental data at *t*_2_ were determined for cells with the expression types N_-_G_-_, N_+_G_-_ and N_-_G_+_ (p < 0.05). No statistically significant differences were found between simulated and experimental data for cells with the expression type N_+_G_+_ at *t*_2_.

The neighbourhood distribution for H1 (see Fig. 1 D) largely agreed with the experimental data at *t*_1_, the only exception being the neighbourhood statistics involving N_-_G_-_ cells. Overall, the neighbourhood distributions fitted the experimental data best, if the cell fate assignment occurred at *t*_0_ = 200 (lowest *Ψ*-value, see Fig. A5). Concerning the neighbourhood structures measured at *t*_2_, the prediction power of the model decreased strongly (see Fig. A4 A).

The neighbourhood statistics for H2 agreed reasonably well with the experimental data (see Fig. 1 E). The neighbourhood pattern of cells with the expression types N_-_G_-_ and N_+_G_-_ at *t*_1_ were similar to the neighbourhood structures obtained under assumption H1, including the misfit involving N_-_G_-_ cells. In addition, simulated N_+_G_+_ cells were significantly less often neighbours of other N_+_G_+_ cells than the experimental data suggests. The latter speaks against a cell fate change of N_+_G_+_ cells into N_-_G_+_ cells.

If performed under the assumption of the H2 model, the evaluated patterns for *t*_2_ showed a better agreement with the experimental data. In particular, the predicted neighbourhood statistics of N_+_G_+_ cells differ statistically less significantly from the *in vitro* measured patterns compared to the simulated cell neighbourhood statistics under the assumption H1. The simulated neighbourhood structures are shown in the Supplementary Material (see. Fig A4B).

According to the overall comparison of neighbourhood structures, the best agreement between experimental data and simulation required *t*_0_ = 200 cells. This implies that all or most cells acquired a cell fate prior to the composition into the organoid, and that after the assembly into a 3D multicellular system, only cell division and cell sorting took place.

### Spatial analysis of expression type distribution

Simulation results of the model hypotheses H1 and H2 fitted well to the experimental data on the spheroid expression type composition, and could also explain the neighbourhood statistics to a large degree. The only disagreement between simulation and experiment concerned the neighbourhood statistics of N_-_G_-_ cells. In order to investigate the reasons for this disagreement, we conducted an additional spatial analysis of the expression type distribution. Fig. 2 shows 3D views of the average ICM organoid composition for both time points *t*_1_ and *t*_2_, marking the spatial density of a given cell fate *j* (see Methods). Cells with the expression type N_+_G_+_ were spread evenly over the whole ICM organoid at *t*_1_ and *t*_2_. The same distribution pattern was obtained for N_+_G_-_ and N_-_G_+_ expression type cells at *t*_1_. Their spatial distribution pattern changed over time. At *t*_2_, N_+_G_-_ cells formed a cluster in the centre of the ICM organoid while N_-_G_+_ cells formed an evenly distributed outer layer around the inner core of the ICM organoid. Concerning the N_-_G_-_ cells, we find that this expression type tended to be positioned in the outer parts of the ICM organoid at both time points, *t*_1_ and *t*_2_. In both cases, their distribution was unevenly spread over the outermost layer of cells in the ICM organoid.

**Figure 2:**
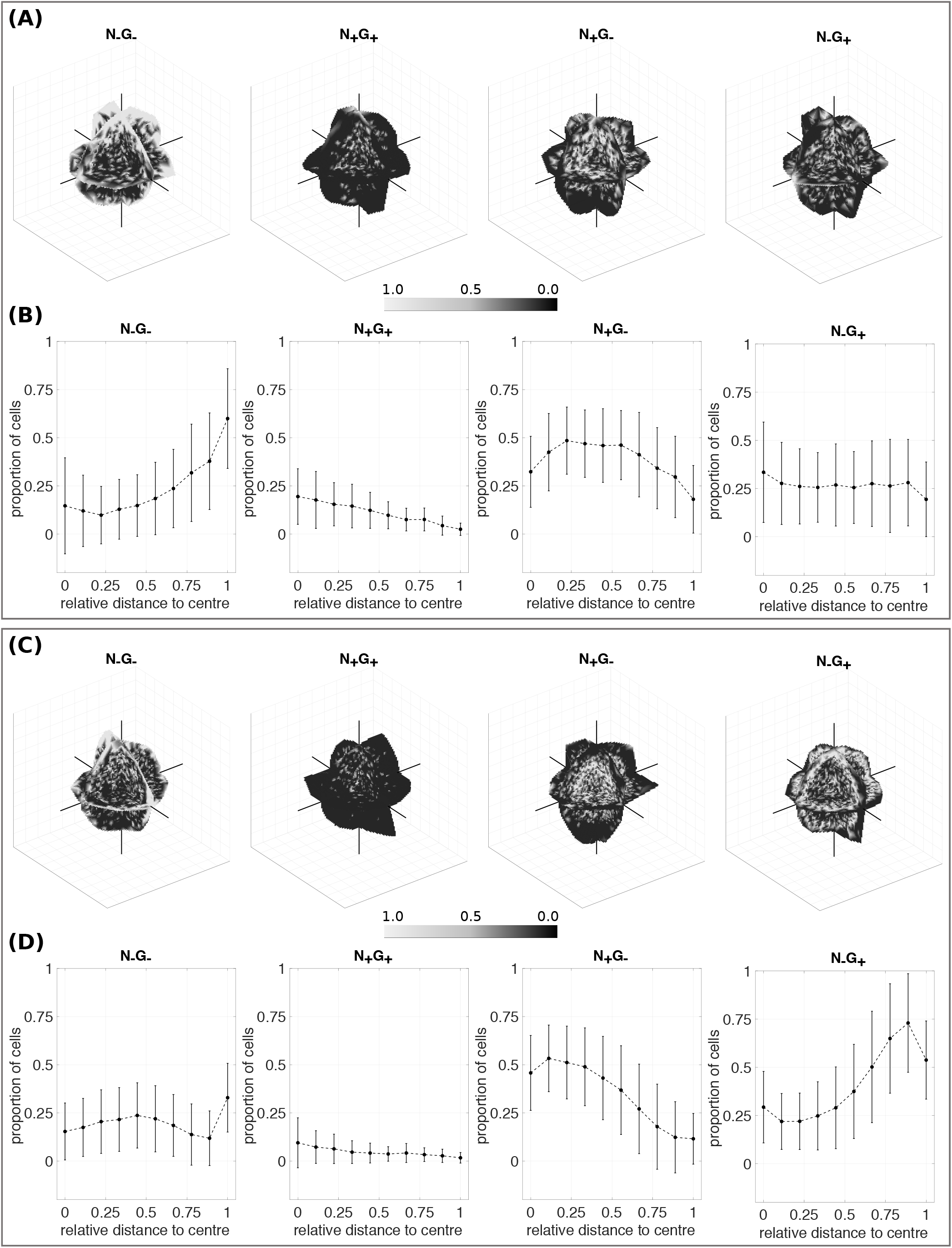
Expression type cluster analysis for ICM organoids. (A) 24 h old ICM organoids; (C) 48 h old ICM organoids. Black indicates the absence of an expression type, white indicates the presence of the selected expression type, respectively. Shown are slices through the ICM organoids at *x* = 0, *y* = 0 and *z* = 0. (B, D) Expression type compositions in dependence of the relative distance to the ICM organoid centre for 24 h old ICM organoids (B) and for 48 h old ICM organoids (D). Cells are sorted according to their distance to the ICM organoid centre of mass and binned into 10 groups. Points indicate the mean proportion of a cell fate type for the 10 bins, the bars denote the standard deviation.

## DISCUSSION

In this study, we investigated whether cell division and cell fate heredity alone can account for cell fate clustering and experimentally observed 3D neighbourhood statistics measured in 24 h and 48 h old ICM organoids, resembling the mid mouse blastocysts (E3.75 to E4.5). Generally, three major processes are hypothesised to lead from undifferentiated ICM cells at E3.0 to the final cell fate arrangement at E4.5 (i.e. inter- and intracellular chemical signalling, cell sorting processes and cell proliferation, reviewed in (20)). We demonstrated that a simple model, involving mechanical interactions such as adhesion and repulsion, cell division with cell fate heredity, and a stochastically driven cell fate, can explain the complex 3D neighbourhood statistics in 24 h old ICM organoids. The proposed model is based on a minimal number of assumptions: (1) Up to a certain time point, when the simulated ICM spheroid consists of a predetermined number of cells (i.e. *t*_0_), all cells coexpress the transcription factors NANOG and GATA6, analogous to mouse embryonic development to E3.0 (11, 45). (2) At *t*_0_, a cell fate is introduced to each cell with a given probability (omitting a detailed description of the signalling pathway dynamics) (45, 46). Cells either continue to co-express both transcription factors (N_+_G_+_), stop the expression of one of them (N_+_G_-_ or N_-_G_+_), or stop to express both (N_-_G_-_). The probabilities for each case are extracted from experimental data (35). (3) Once a certain cell fate has been chosen, cells pass on their cell fate to the daughter cells during cell division according to certain rules (see Fig. A1). Four hypotheses addressing cell fate switches during cell division were tested: The cell fate is passed on to both daughter cells, switches between different fates are not allowed (H1); N_+_G_+_ cells give rise to N_+_G_+_ or N_-_G_+_ cells (H2); N_+_G_+_ cells give rise to N_+_G_+_, N_+_G_-_ or N_-_G_+_ cells and N_+_G_-_ can switch to N_+_G_-_ (H3); H3 is extended, such that N_+_G_-_ cells can switch to N_+_G_-_ and vice versa (H4).

### Proportions

Since the initial cell fate assignment at different *t*_0_ was adjusted to fit the proportion data known from *t*_1_, all models reproduced the proportions for *t*_1_ very well.

At *t*_2_, N_+_G_-_ cells represented the majority in the ICM organoid. The increase of N_-_G_+_ cells from *t*_1_ to *t*_2_ can be explained by either cell fate switches from N_+_G_-_ to N_-_G_+_ or by an enhanced cell division rate for N_-_G_+_ cells, for instance due to an interaction between the cell growth rate and mechanical forces exerted by surrounding cells. The first can be considered as unlikely, regarding the overall fit of the simulated patterns to experimental data (see Fig. A3). These results indicate that cell fate switches from N_+_G_-_ to N_-_G_+_ and vice versa are unlikely, which is in agreement with the literature (16, 17, 19).

Cells on the surface of ICM organoids might grow faster than cells in the centre of ICM organoids, where the cell density is high (37, 47). N_-_G_+_ expressing cells are predominantly found at the rim of ICM organoids where the cell density is lower, which might result in a higher cell division rate (24, 48). A feedback of pressure onto cell growth is not included in our model, hence this aspect has not been captured.

### Neighbourhood structures

Complex 3D spatial arrangements of cells with particular cell fates characterise early stages of PrE and Epi segregation. ICM organoids mimic those cell arrangements after 24 h of growth and after 48 h of growth for mid and late mouse blastocysts, respectively. Mathew *et al.* (2019) (35) used a computational rule-based static model in order to reproduce the cell fate pattern measured in 24 h old ICM organoids. Hereby, four different hypotheses have been tested. The first three simulations were based on hypotheses derived from the salt-and-pepper pattern, while the fourth tested pattern was based on a local density. Patterns from all four simulations were significantly different from experimentally observed patterns (35), in particular because the clustering of the cells sharing the same cell fate could not be reconciled with any of the tested models.

In this study, we showed that an initial salt-and-pepper distributed cell fate decision, in addition to cell division involving cell fate heredity, adhesion and mechanical repulsion, can explain the lineage composition and spatial distribution of N_+_G_-_ and N_+_G_-_ cells in 24 h old ICM organoids. The main assumption here was that the 24 h time point did not represent the time point of the cell fate decision. Instead, we assumed that the cell fate decision occurred previously, and that the local clustering of cell fates arises from cell divisions. Under this hypothesis, we found that the assumption H1 as well as the assumption H2 showed good agreement with experimental data for 24 h old ICM organoids, whereby the assumption H1 reflected the neighbourhood structure of N_+_G_+_ cells better than the assumption H2. With increasing complexity of the cell fate heredity rules, the overall goodness of the fit of the simulated compared to the experimentally determined neighbourhood statistics decreases. Hence, the experimental data is explained best, if the cell fates are determined at *t*_0_, and a phase of cell divisions follows.

A comparable model, which is also based on mechanic cell displacement and cell division, but takes also into account an intercellular network for cell fate decisions was used in (41) to predict cell fate decisions in *in vivo* mouse blastocysts. According to their findings, the spatio-temporal distribution of PrE and Epi cells follows the salt-and-pepper pattern, whereby Epi cells are preferentially surrounded by PrE cells and vice-versa (41).

The distribution of cell fate patterns is questioned by several recent studies that are highlighting the need of neighbourhood interactions for the rise of Epi and PrE lineages (25, 27, 41). However, we hypothesise that the local spatial clustering of cell fates as obtained in (35) might not be indicative of specific neighbourhood interactions involved in cell fate decision. We further hypothesise that a cell fate clustering is a transient state, characteristic of blastocysts and ICM organoids in the time span between cell fate decision and spatial sorting. In particular, our results indicate that an initial cell fate decision based on a random distribution, in addition to cell divisions and cell fate heredity, can give rise to local cell clusters, exhibiting the same neighbourhood statistics as observed in experiments (35).

With the given restrictions on the cell fate switches, it was not possible to fit the proportion transition from 24 h to 48 h data perfectly well. Although cell fate switches are considered unlikely, we investigated if the proportion data and neighbourhood structures could be approximated better if further cell fate switches are allowed. Indeed, this relaxation allows for a better fit of the cell proportion data. However, the neighbourhood statistics are not approximated well under the hypotheses H3 and H4 (see Supplementary Material).

Using the model, we can provide an estimate for the time point when the cell fate decision process in ICM organoids is finalised. Since the simulated data fits best to the experimental data when the cell fate is introduced at a cell count of 200 cells, we assume that *in vitro*, most cells of the ICM organoids acquired a cell fate after the composition into the organoid and before their first cell division.

This implies that the initial cell fate acquisition is a process which takes place directly after the ICM organoid composition from single cells. Propagating in time, cell division leads to a local clustering of cell fates and a cell sorting process yields the final separation of PrE and Epi cells. Krupinski *et al.* (2011) hypothesised differential adhesion between cells of different fates as a critical determinant in the robust ICM formation (49).

### Spatial analysis of expression type distribution

NANOG and GATA6 are described as the first markers for Epi and PrE cell fate segregation, with a salt-and-pepper like occurrence in early and mid mouse blastocyst. During blastocyst growth, PrE cells are sorted to the rim of the ICM, while Epi cells remain in the centre of the ICM (1, 7, 10–17). This behaviour of PrE and Epi cells is reflected in ICM organoids. While N_+_G_-_ and N_-_G_+_ cells are evenly distributed over the 24 h old ICM organoid, they re-localise in 48 h old ICM organoids, forming a centre of N_+_G_-_ cells and a rim of N_-_G_+_ cells. N_+_G_+_ cells are co-expressing NANOG and GATA6, as described for early mouse blastocysts (E3.0) (7, 11, 12). Thus, we expected them to be distributed evenly over the whole ICM for 24 h old and 48 h old ICM organoids, which was confirmed by the conducted spatial analysis. For N_-_G_-_ cells, expressing low levels of NANOG and GATA6, we also expected a spatially homogeneous distribution. However, the spatial analysis revealed that N_-_G_-_ cells tended to be positioned at the rim of ICM organoids at the 24 h time point. This finding was surprising, but it explains the very high proportion of N_-_G_-_ cells in the neighbourhood of N_-_G_-_ cells. Our model assumptions treat all cell fates equally and do not involve a specific spatial positioning of one of the cell fates. Furthermore, we assume that the original cell fate distribution follows a salt-and-pepper pattern (following from the experimental procedure to compose the initial organoids). Apparently, a subgroup of the N_-_G_-_ cells tends to assemble at the rim of the organoid. This behaviour has not been described in the literature before and is not yet understood. However, this positioning explains the disagreement between the expected neighbourhood statistics and the data for this cell type. The role and dynamics of N_-_G_-_ cells is not yet well understood, and should be investigated in further studies.

### Conclusions

With this study we aimed to clarify the contributions of cell proliferation to the observed neighbourhood statistics in ICM organoids, neglecting intra- and intercellular chemical signalling as well as a cell sorting process. Assuming an initially random cell fate assignment, subsequently followed by cell divisions and cell displacement in response to mechanical cell-cell interactions, the *in vitro* observed neighbourhood statistics can be reproduced reasonably well.

The agreement between simulated and experimental data is best for the simplest assumptions for cell fate changes during cell division (i.e. cells strictly pass on their cell fate to daughter cells). Although the simulated neighbourhood structures fit parts of the experimental data reasonably well, the model is lacking a cell sorting process, which we hypothesise to result in the measured neighbourhood structures. Cells placed at the rim of cell colonies are described to grow faster, and thus show a higher cell division rate (37, 47). After the sorting of GATA6 positive cells to the rim of the ICM organoid, these cells might divide with a higher rate, which would explain the strong increase of their proportions from 24 h to 48 h old ICM organoids.

The results of our study indicate a temporal separation of the four phases shown in Fig. 3. Further indication for this assumption can be found in the work of Filimonow *et al.* 2019 (50). Here it has been shown that no difference in the E-cadherin levels between Epi and PrE can be found until E3.75. This implies that cell sorting does not take place until this time point, if cell sorting partially occurs due to differential adhesion (49, 51, 52).

**Figure 3:**
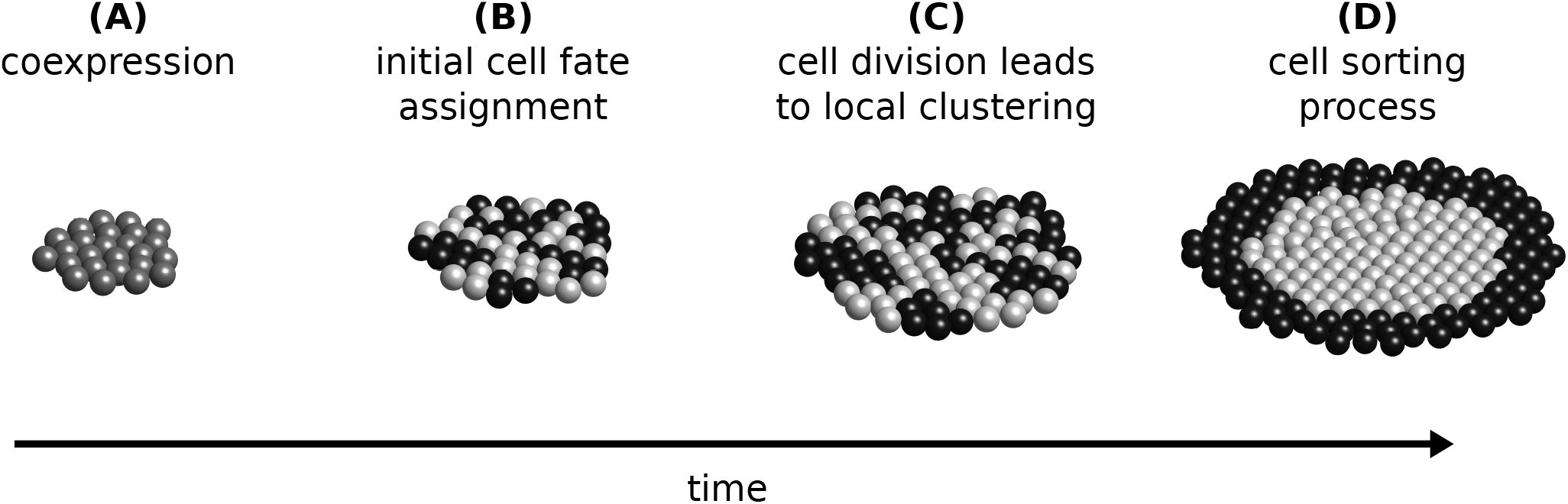
Working model. Over time, four phases lead to the separation of GATA6 positive (black) and NANOG positive (white) cells. Initially, all cells coexpress both cell fate markers (A), followed by a random cell fate assignment (B). Cell division leads to a local clustering of cells sharing the same cell fate (C) and eventually a sorting mechanism leads to a phase separation of GATA6 positive and NANOG positive cells.

In particular, our results indicate that the cell fate decision takes place prior or only shortly after the organoids are assembled from 200 cells, and that the neighbourhood structure between the time points *t*_0_ and *t*_1_ are determined only by cell divisions and cell fate heredity. Subsequently, cell sorting starts after *t*_1_, which is supported by (50), and eventually leads to the phase separation found 48 h old ICM organoids.

## AUTHOR CONTRIBUTIONS

Conception and design of the study were performed by T.L. and F.M.; Experimental data were provided by B.M., S.C.F. and E.H.K.S.; T.L. implemented the code.; T.L., A.D. and F.M. analysed the data; T.L., F.M. and S.C.F. interpreted the data. T.L. wrote the manuscript. All authors revised the manuscript. F.M. supervised the study.

## ACKNOWLEDGMENTS

Research in the F.M. lab is supported by the Giersch foundation. T.L., F.M. and E.H.K.S. are supported by a grant from the Hessen State Ministry for Higher Education, Research and the Arts in the framework of the Loewe Program (DynaMem). Research in the Stelzer lab is supported by the Deutsche Forschungsgemeinschaft (CEF-MC II, EXC-115). B.M. has additionally been supported by fellowships from the Joachim Herz Stiftung and Freunde und Förderer der Goethe-Universität. S.C.F. and E.H.K.S. have been supported by an International Exchanges Grant from The Royal Society (IE141022).

