## Supplementals for "Cell Fate Clusters in ICM Organoids Arise from Cell Fate Heredity & Division – a Modelling Approach"

### SUPPLEMENTARY MATERIAL

#### Parameter estimation for different hypotheses

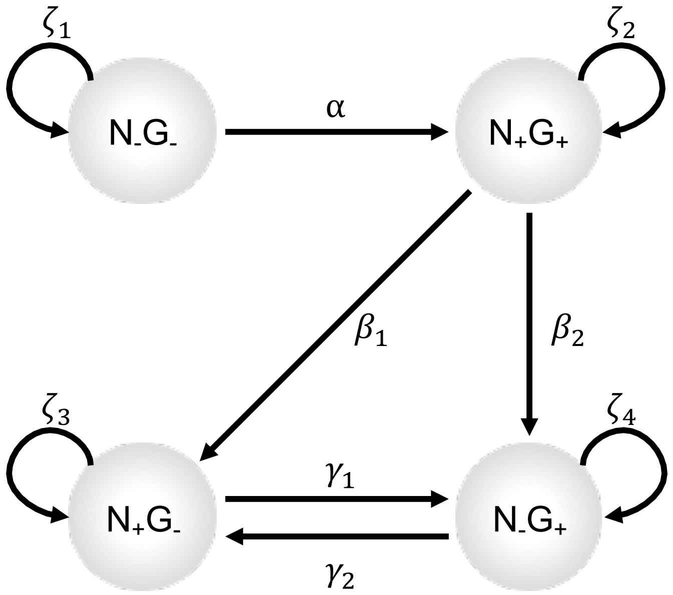


Figure A1: Hypotheses for cell fate switches during cell division. The following cell fate switches are considered: $N_-G_-$ remain $N_-G_-$ or become $N_+G_+$. $N_+G_+$ remain $N_+G_+$ or become $N_+G_-$ or $N_-G_+$. $N_+G_-$ and $N_-G_+$ remain $N_+G_-$ and $N_-G_+$ or switch to the opposite cell fate, respectively. The probabilities for the different cell fate transitions distinguish the different models (see Tab. A2).

The model can be used to test several assumptions addressing cell fate heredity. In total, four different hypotheses are tested. Each hypothesis is considering cell fate switches during cell division according to Fig. A1. During cell division, the cell fate is passed on to the daughter cells. A cell fate switch is possible with a given rate. The model considers the following cell fate switches: $N_-G_-$ remain $N_-G_-$ ($\zeta_1$) or become $N_+G_+$ ($\alpha$). $N_+G_+$ remain $N_+G_+$ ($\zeta_2$) or become $N_+G_-$ ($\beta_1$) or $N_-G_+$ ($\beta_2$). $N_+G_-$ and $N_-G_+$ remain $N_+G_-$ ($\zeta_3$) and $N_-G_+$ ($\zeta_4$) or switch to the opposite cell fate ($\gamma_1$) and ($\gamma_2$), respectively (see Fig. A1). These cell fate transitions form a system of linear ordinary differential equations, which can be written as:

$$d\mathbf{f}/dt = \mathbf{A}\mathbf{f} \tag{A1}$$

$$\begin{pmatrix} dN_+G_+/dt \\ dN_-G_-/dt \\ dN_+G_-/dt \\ dN_-G_+/dt \end{pmatrix} = \begin{pmatrix} \zeta_2 - \beta_1 - \beta_2 & \alpha & 0 & 0 \\ 0 & \zeta_1 - \alpha & 0 & 0 \\ \beta_1 & 0 & \zeta_3 - \gamma_1 & \gamma_2 \\ \beta_2 & 0 & \gamma_1 & \zeta_4 - \gamma_2 \end{pmatrix} \cdot \begin{pmatrix} N_+G_+ \\ N_-G_- \\ N_+G_- \\ N_-G_+ \end{pmatrix}, \tag{A2}$$

with the analytical solution:

$$\mathbf{f}(t) = c_{N_+G_+}\mathbf{v}_{N_+G_+}e^{\lambda_{N_+G_+}t} + c_{N_-G_-}\mathbf{v}_{N_-G_-}e^{\lambda_{N_-G_-}t} + c_{N_+G_-}\mathbf{v}_{N_+G_-}e^{\lambda_{N_+G_-}t} + c_{N_-G_+}\mathbf{v}_{N_-G_+}e^{\lambda_{N_-G_+}t}, \tag{A3}$$

with $\mathbf{v}$ and $\lambda$ the eigenvectors and eigenvalues of the coefficient-matrix $\mathbf{A}$, respectively. The unknown $c$ values can be determined by inserting the known cell counts of the different cell types at $t_1$. The rates for the different cell fate switches vary for the different hypotheses (see Tab. A2).

Table A1: Chosen values for model parameters.

| **Parameter** | **Value** |
|---|---|
| $F_0$ | 1 |
| $a$ | 0.6 |
| $\sigma$ | 4 |
| $k$ | 1 |
| $r^*$ | 1 |

Table A2: Cell fate transition rates for the four different hypotheses.

| | **hypothesis** | | | |
|---|---|---|---|---|
| **parameter** | **H1** | **H2** | **H3** | **H4** |
| $\zeta_1$ | 1 | 1 | 1 | 1 |
| $\zeta_2$ | 1 | 1 | 1 | 1 |
| $\zeta_3$ | 1 | 1 | 1 | 1 |
| $\zeta_4$ | 1 | 1 | 1 | 1 |
| $\alpha$ | 0 | 0 | 0.2123 | 0.2123 |
| $\beta_1$ | 0 | 0 | 0.7863 | 0.7863 |
| $\beta_2$ | 0 | 1 | 0.7863 | 0.7863 |
| $\gamma_1$ | 0 | 0 | 0.2997 | 0.3927 |
| $\gamma_2$ | 0 | 0 | 0 | 0.1 |

**Flowchart**

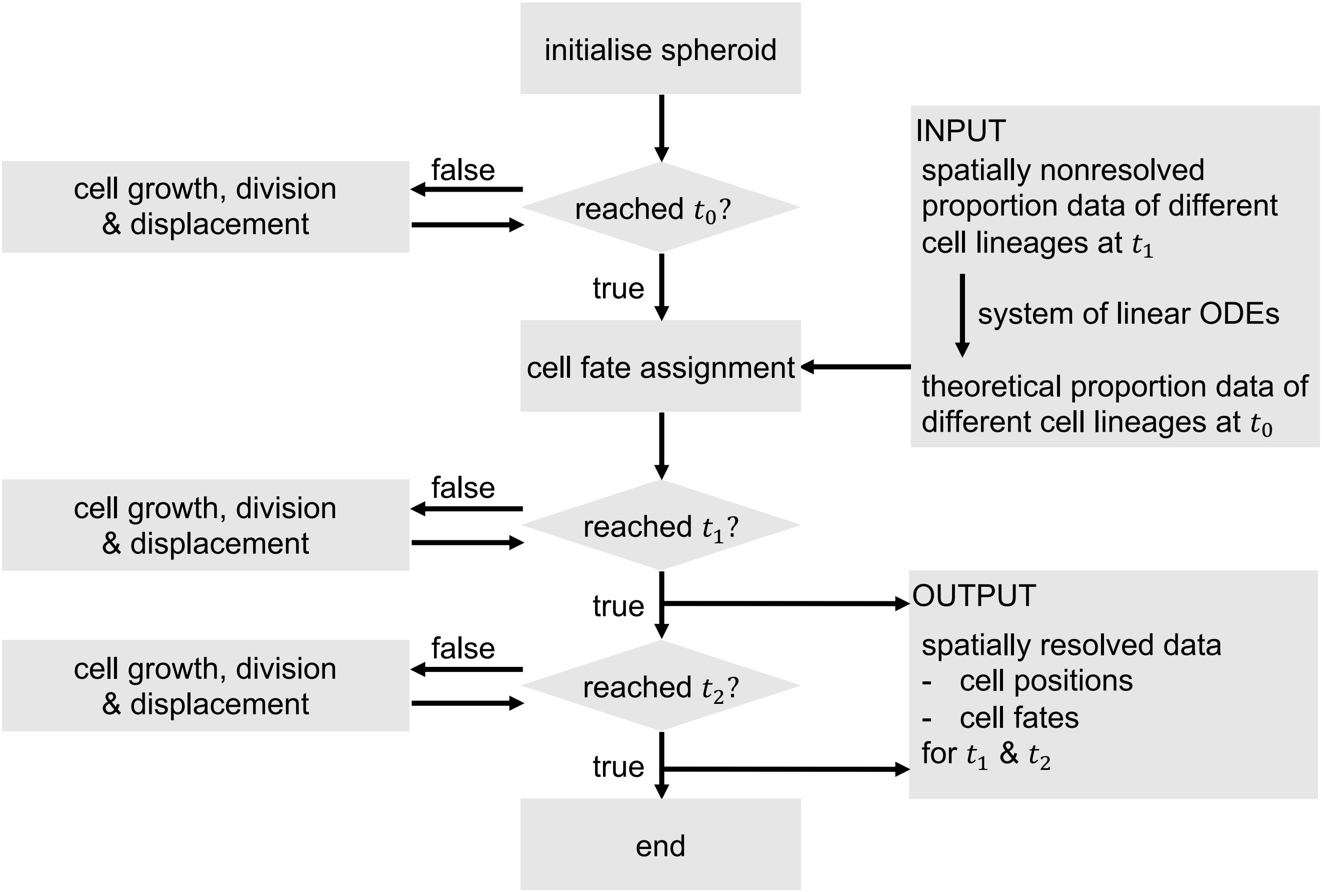


Figure A2: Flowchart of the implemented model.

### Hypothesis 3 and 4

If we consider cell fate switches from $N_+G_-$ to $N_-G_+$ and vice versa, which according to the literature are considered as unlikely, a third and a fourth hypothesis can be formulated. In both hypotheses, the strong increase in the amount of $N_-G_+$ is explained by cell fate switches from $N_+G_-$ to $N_-G_+$. While H3 permits only the cell fate switches from $N_+G_-$ to $N_-G_+$, H4 is considering a small flux between both cell fates. The parameter values for both hypotheses are shown in Tab. A2.

As expected, the simulated proportions for H3 agree very well with the experimental data at $t_1$ and $t_2$ (see. Fig. A3 A and B).

Comparing H3 to H1 and H2, we observe that its $\psi$ values are higher, thus the goodness of the fit of the neighbourhood statistics is lower (see. Fig. A3 B). The neighbourhood distributions largely agree with experimental data at $t_1$ for the neighbourhood of double positive and NANOG positive cells. The simulated proportions of GATA6 positive cells adjacent to other GATA6 positive cells are significantly lower compared to experimental data, independent of the cell count of the ICM spheroid at which the initial cell fate is determined (see. Fig. A3 C).

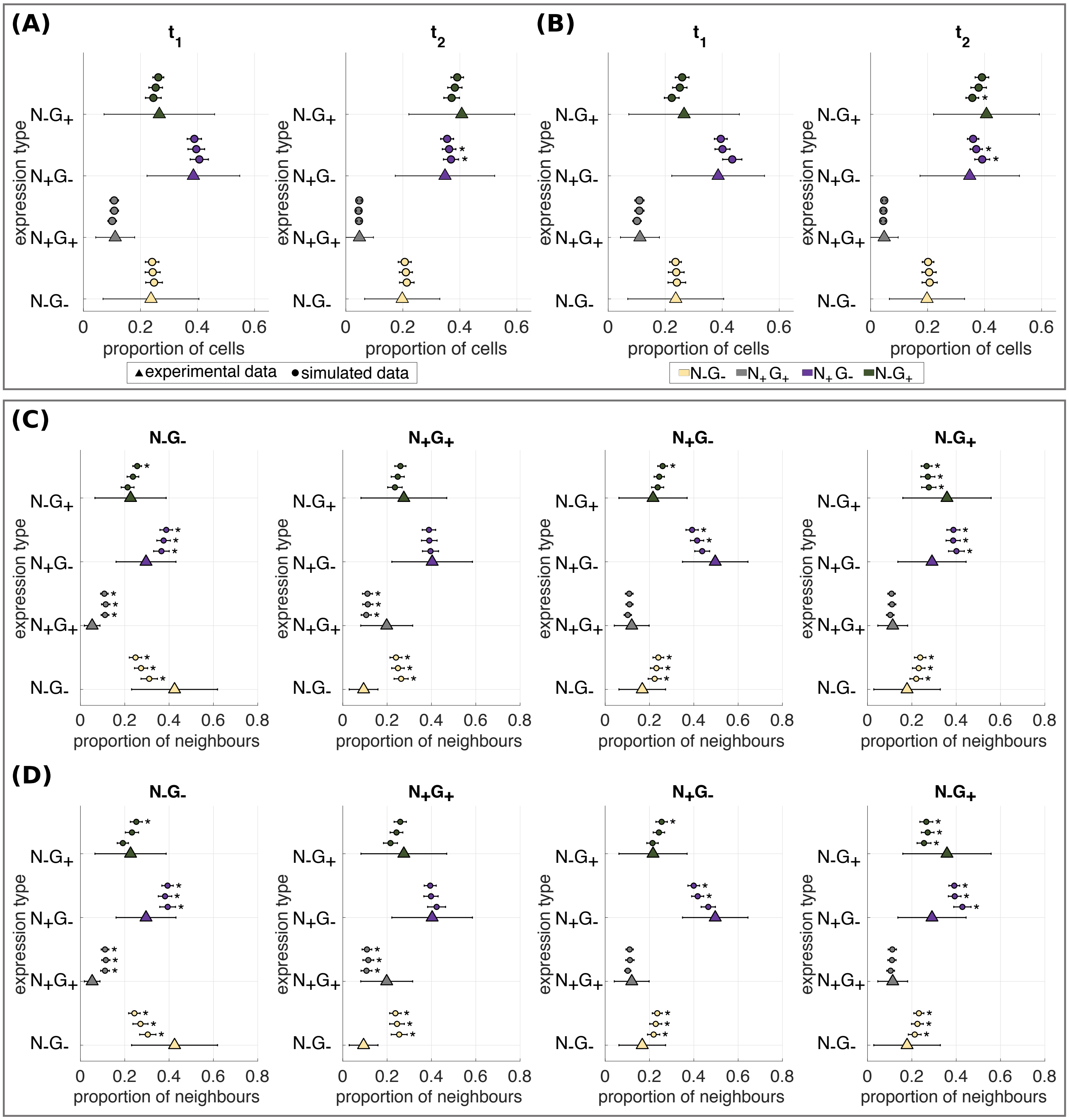


Figure A3: Expression type composition of ICM spheroids for H3 and H4. (A, B) Expression type composition of ICM organoids and ICM spheroids shown as percentage of the total number of cells within ICM organoids at $t_1$ and $t_2$. Statistically significant differences between the cell fate proportion of ICM organoids and ICM spheroids are indicated by stars ($p < 0.05$; using a Wilcoxon-Mann-Whitney test with Bonferroni correction). Simulations were performed under the assumption H3 (A) and the assumption H4 (B). (C, D) Expression type composition of neighbouring cells shown as percentage of the total of neighbouring cells at $t_1$. Simulations were performed under the assumption H3 (C) and the assumption H4 (D). Experimental data from Mathew *et al.* (2019) are indicated by triangles. Simulation results for different $t_0$ are indicated by circles. The error bars indicate the standard deviation. $t_0$ from lowest line to top: 200, 300 and 400 cells. Statistically significant differences between the neighbourhood structure of 24 h old ICM organoids and ICM spheroid patterns are indicated by stars ($p < 0.05$; using a Wilcoxon-Mann-Whitney test with Bonferroni correction).

### Neighbourhood structures at $t_2$

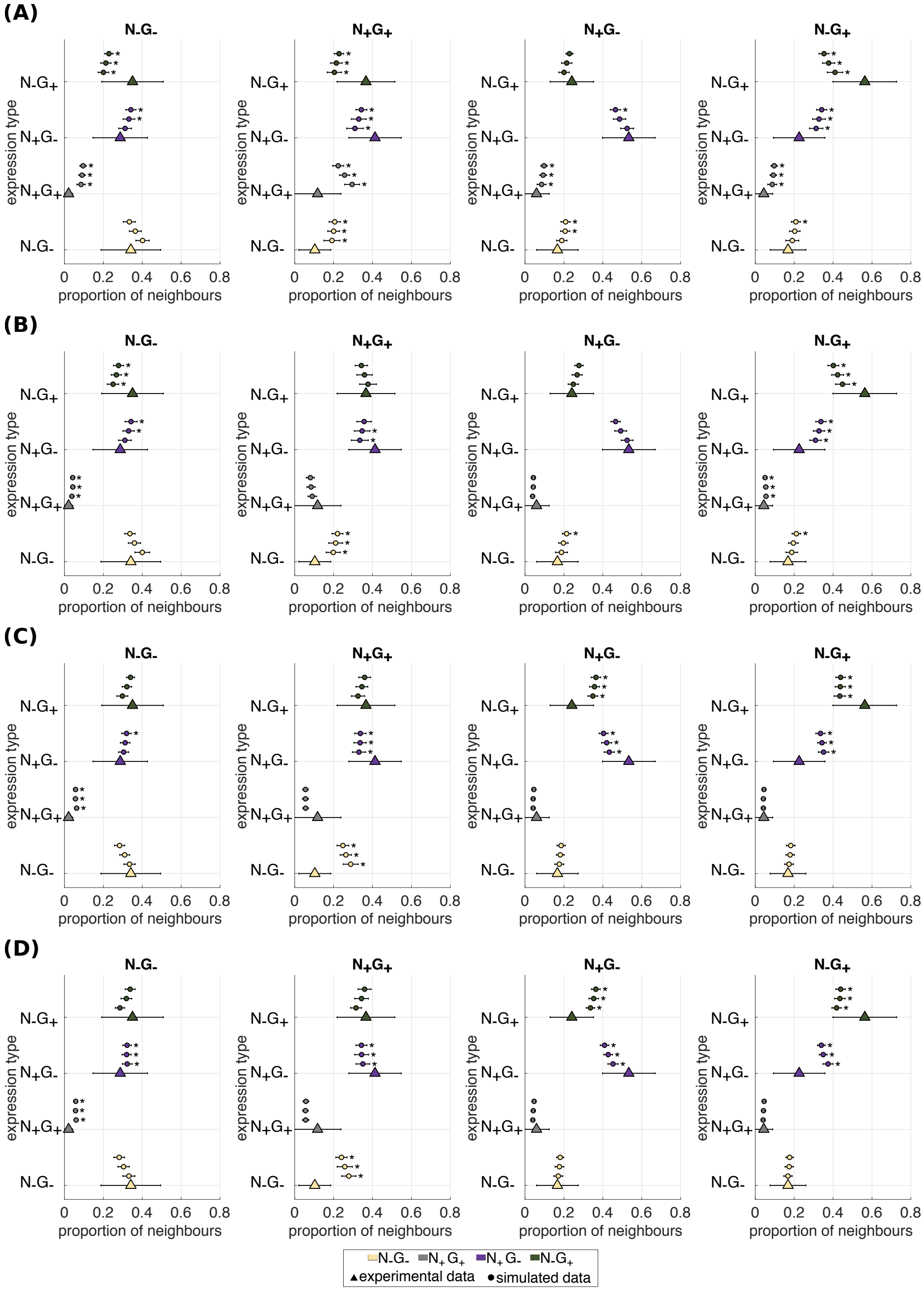


Figure A4

Figure A4: Expression type composition of neighbouring cells shown as percentage of the total of neighbouring cells at $t_2$ for H1-H4. Simulations were performed under the assumption H1 (A), the assumption H2 (B), the assumption H3 (C), and the assumption H4 (D). Experimental data from Mathew *et al.* (2019) are indicated by triangles. Simulation results for different $t_0$ are indicated by circles. The error bars indicate the standard deviation. $t_0$ from lowest line to top: 200, 300 and 400 cells. Statistically significant differences between the neighbourhood structure of 24 h old ICM organoids and ICM spheroid patterns are indicated by stars ($p < 0.05$; using a Wilcoxon-Mann-Whitney test with Bonferroni correction).

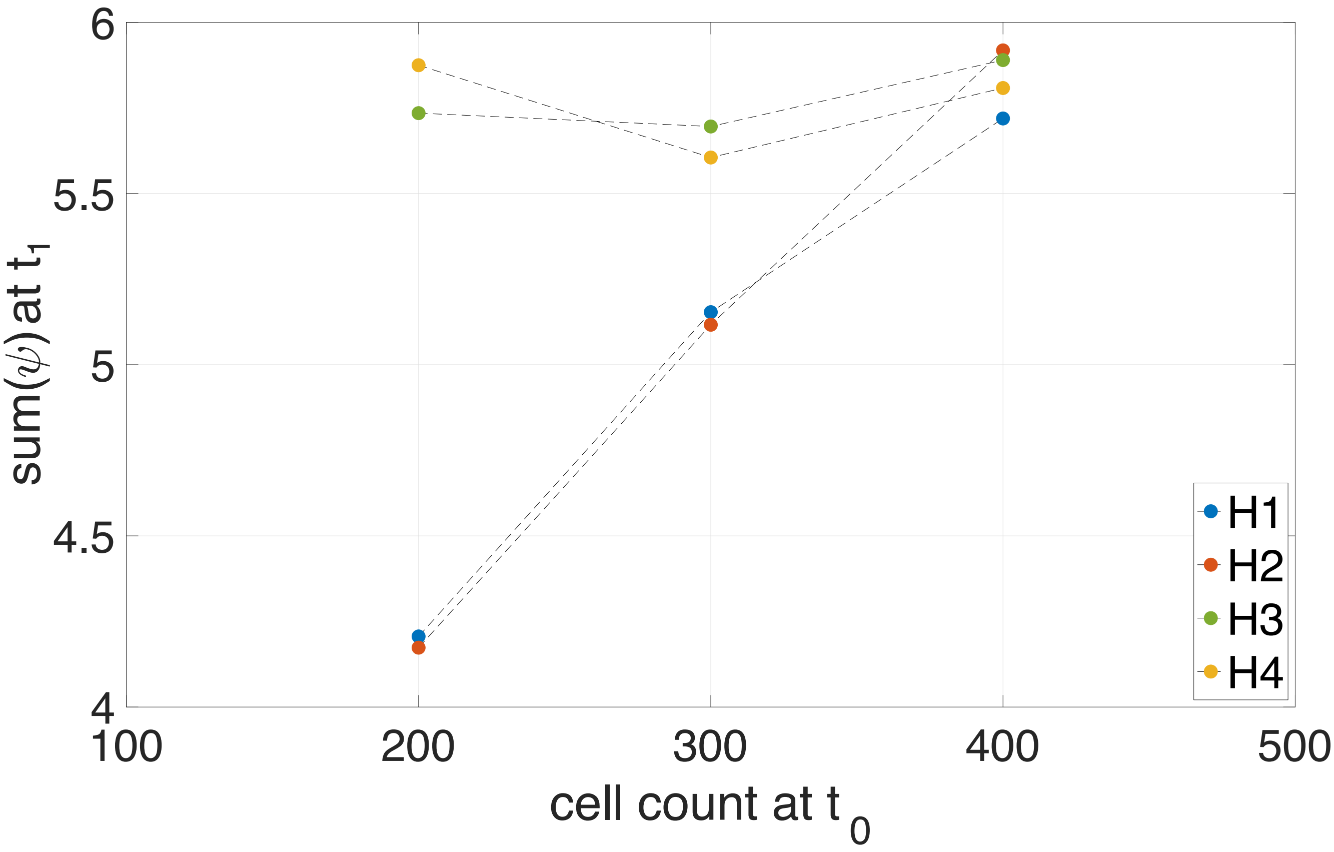


Figure A5: Sum of all $\psi$ values at $t_1$ is shown for H1, H2, H3 and H4.
